# Reservoir-based Tracking (TRAKR) For One-shot Classification Of Neural Time-series Patterns

**DOI:** 10.1101/2021.10.13.464288

**Authors:** Muhammad Furqan Afzal, Christian David Márton, Erin L. Rich, Helen Mayberg, Kanaka Rajan

**Author notes:** These authors contributed equally to this work (Muhammad Furqan Afzal, Christian David Márton).

## Abstract

Distinguishing between complex nonlinear neural time-series patterns is a challenging problem in neuroscience. Accurately classifying different patterns could be useful for a wide variety of applications, e.g. detecting seizures in epilepsy and optimizing control spaces for brain-machine interfaces. It remains challenging to correctly distinguish nonlinear time-series patterns because of the high intrinsic dimensionality of such data, making accurate inference of state changes (for intervention or control) difficult. On the one hand, simple distance metrics, which can be computed quickly, often do not yield accurate classifications; on the other hand, ensembles or deep supervised approaches offer high accuracy but are training data intensive. We introduce a reservoir-based tool, state tracker (TRAKR), which provides the high accuracy of ensembles or deep supervised methods while preserving the benefits of simple distance metrics in being applicable to single examples of training data (one-shot classification). We show that TRAKR instantaneously detects deviations in dynamics as they occur through time, and can distinguish between up to 40 patterns from different chaotic data recurrent neural networks (RNNs) with above-chance accuracy. We apply TRAKR to a benchmark time-series dataset – permuted sequential MNIST – and show that it achieves high accuracy, performing on par with deep supervised networks and outperforming other distance-metric based approaches. We also apply TRAKR to electrocorticography (ECoG) data from the macaque orbitofrontal cortex (OFC) and, similarly, find that TRAKR performs on par with deep supervised networks, and more accurately than commonly used approaches such as Dynamic Time Warping (DTW). Altogether, TRAKR allows for high accuracy classification of time-series patterns from a range of different biological and non-biological datasets based on single training examples. These results demonstrate that TRAKR could be a viable alternative in the analysis of time-series data, offering the potential to generate new insights into the information encoded in neural circuits from single-trial data.

## 1 Introduction

The size and complexity of neural data collected has increased greatly ([22]). Neural data display rich dynamics in the firing patterns of neurons across time, resulting from the recurrently connected circuitry in the brain. As our ability to record neural dynamics increases through new technologies, so does the need to understand how these dynamical patterns change across time and how they relate to one another.

A lot of work in computational neuroscience over the past decade has focused on modeling the collective dynamics of a population of neurons in order to gain insight into how firing patterns are related to task variables ([23, 30, 46, 28, 17, 47, 24, 9, 8, 26, 39, 21, 38, 4, 37]). These approaches, however, rely on fitting the whole dynamical system through many rounds of optimization with multiple batches of training data (or multiple “trials”), either indirectly–by modeling the task inputs and outputs ([23, 17, 8, 39, 21, 38]), or directly–by adjusting the weights of a neural network to fit recorded firing patterns ([24, 9]). The reliance on multiple trials makes it difficult to use such approaches to glean insights from single-trial data, or in a regime where recordings cannot be repeated easily across multiple trials. This problem arises in a clinical setting, for example in the detection of seizures or during deep brain stimulation where patterns need to be assessed in a real-time, single-session setting.

Previous approaches for classifying time series lie on a spectrum from simple distance metrics (e.g., Euclidean) to more computationally intensive approaches such as Dynamic Time Warping (DTW) ([45]), ensembles of classifiers ([3]) or deep supervised learning ([15, 10]). On one end, computing simple distance metrics is fast and straightforward, but does not always yield high accuracy results because the patterns may not be perfectly aligned in time. On the other end of the spectrum, ensembles of classifiers and deep learning based approaches ([3, 15, 10]) have been developed that can offer high accuracy results, but at high computational cost (leveraging multiple rounds of optimization and batches of training data). DTW, in the middle of the spectrum, has been found to offer good results relative to computational cost ([10, 2, 35]) and is now routinely used to measure the similarity of time-series patterns. Altogether, though, there are few approaches that offer high accuracy results at relatively low computational cost, which can also be used on single examples of training data, such as from single trials or single sessions.

An alternative could be found in reservoir computing, in which prior work has shown that networks of neurons can be used as “reservoirs” of useful dynamics, so called echo-state networks (ESNs), without the need to train recurrent weights through successive rounds of expensive optimization ([42, 25, 41, 6, 14, 13, 12, 20]). This suggests reservoir networks could offer a computationally cheaper, yet viable, alternative to deep supervised approaches in the classification of neural time-series data. An example is the time-warping invariant echo-state network (twiESN) which can handle time-warped signals for classification and performs comparably to DTW ([40]). However, the training of reservoir networks has been found to be more unstable compared to methods that also adjust the recurrent connections (e.g., via backpropagation through time, BPTT) in the case of reduced-order data ([42]). Even though ESNs have been shown to yield good results when fine-tuned ([40, 1]), convergence represents a significant problem when training ESNs end-to-end to perform classification on complex time-series datasets and is a hurdle to their wider adoption. We offer a way around these problems here.

Addressing the need for high performing approaches at low computational cost and taking inspiration from reservoir computing, we propose *state tracker* (TRAKR for short). With TRAKR, we fit the readout weights from a reservoir to a *single* time series – thus avoiding many rounds of optimization that increase training time and can cause instabilities. We then freeze the readout weights, and compute the error between the network output, generated in response to a particular test pattern, and its target (Figure 1).

**Figure 1:**
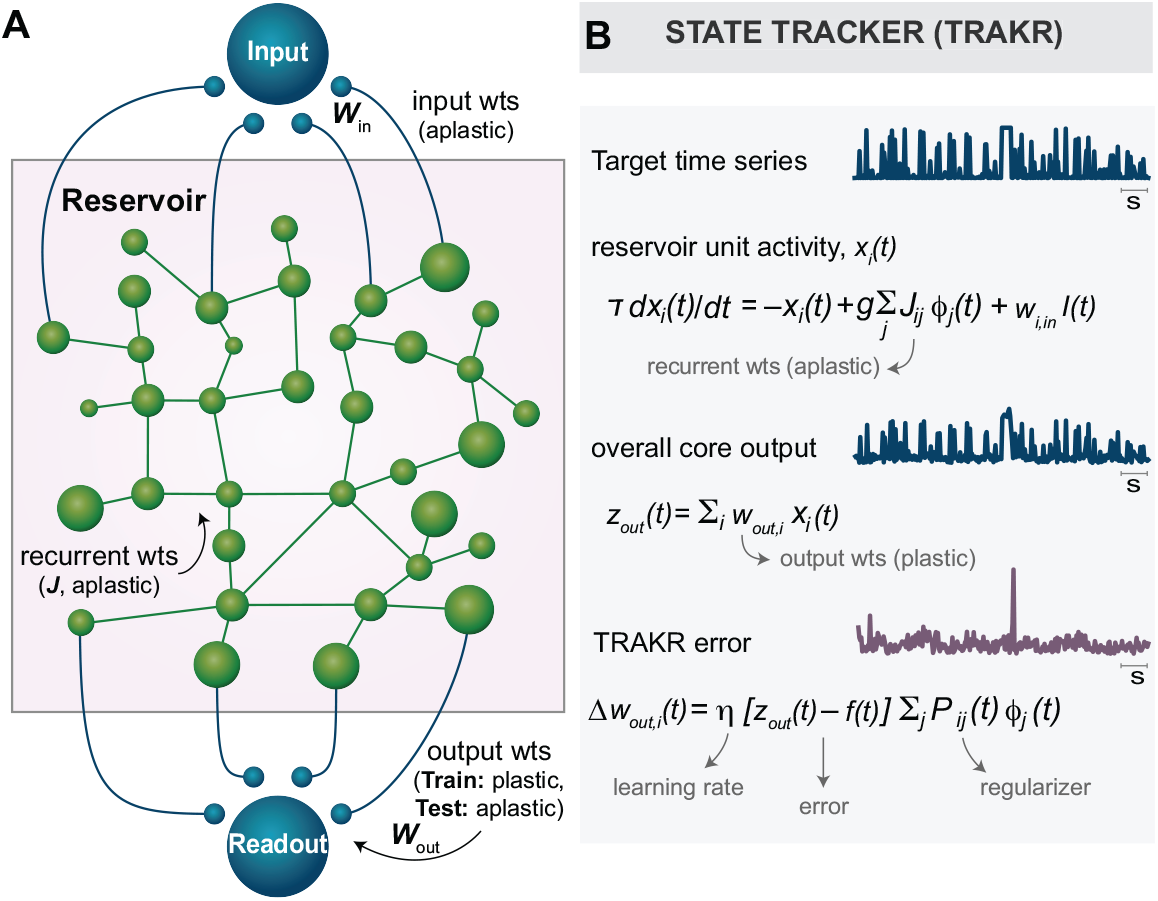
A) TRAKR setup overview. TRAKR consist of a reservoir connected to input and readout units via dedicated weights. Recurrent weights J and input weights *w_in_* are aplastic. Only the output weights *w_out_* are subject to training. B) TRAKR equations for single unit activity, readout unit activity and error term.

We show that using TRAKR, time-series patterns generated from ≈ 10 chaotic data recurrent neural networks (RNNs) could be distinguished with 90% accuracy, and up to ≈ 40 patterns with above-chance accuracy. We also obtained high accuracy results on a benchmark dataset–permuted sequential MNIST–outperforming other approaches such as simple distance metrics and a previous approach based on echo-state networks, while performing on par with supervised approaches leveraging deep neural networks.

We also tested TRAKR on neural data from the macaque orbitofrontal cortex (OFC), a higher-order brain region involved in encoding expectations and inducing behavioral changes to unexpected outcomes ([29, 31, 16, 43, 32, 7, 44, 33]). We fitted TRAKR to single-trial data from the rest period and probed whether it could distinguish between the neural signals from 3 different behaviorally-relevant periods–(rest, *choice*, *reward seeking*). We found that TRAKR was able to distinguish the three epochs with high accuracy, showing that it can be used as a reliable tool to assess the similarity of patterns occurring through time in neural circuits based on single trial data.

We also applied TRAKR to another neural dataset, 8-channel data recorded from the subcallosal cingulate (SCC) region of human patients with treatment-resistant depression who are undergoing deep brain stimulation in the operating room (OR) ([34, 36]). We found that TRAKR-derived reservoir activations can be used to better distinguish the effect of stimulation on neural patterns, achieving greater separation in latent space compared to raw traces. This shows how latent representations can be visualized for use in the clinical domain, and points to the ability of TRAKR to distinguish complex neural time-series patterns.

Taken together, we make the following contributions:

- We show that we can use reservoirs to derive an error signal from single-trial data which can be used to reliably distinguish time-series patterns
- We show that our approach, TRAKR, performs better than routinely used approaches such as distance metrics (e.g. Euclidean distance, or Mutual Information) and Dynamic Time Warping (DTW), and on par with computationally more intensive approaches based on multiple batches of training data such as supervised approaches based on deep neural networks
- We show that TRAKR achieves high accuracy results on a benchmark dataset, permuted sequential MNIST
- We show that TRAKR also achieves high accuracy results on biological data, where it can be used to distinguish complex time-series patterns

Altogether, due to its high accuracy and computationally light profile, TRAKR is an attractive option for generating insights into the information encoded in neural signals based on single-trial data.

## 2 Methods

### 2.1 Model details

TRAKR (Figure 1) is a reservoir-based recurrent neural network (RNN) with *N* recurrently connected neurons. Recurrent weights, *J*, are initialized randomly and remain aplastic over time ([6, 12, 20]). The readout unit, *z_out_*, is connected to the reservoir through a set of output weights, *w_out_*, which are plastic and are adjusted during training. The reservoir also receives an input signal, *I*(*t*), through an aplastic set of weights *w_in_*.

The network is governed by the following equations:

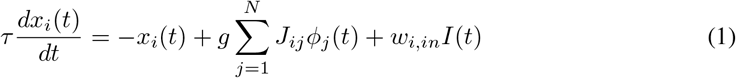

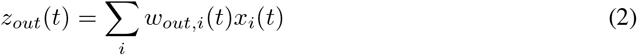

Here, *x_i_*(*t*) is the activity of a single neuron in the reservoir, *τ* is the integration time constant, *g* is the gain setting the scale for the recurrent weights, and *J* is the recurrent weight matrix of the reservoir.

The term 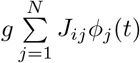 denotes the strength of input to a particular neuron from other neurons in the reservoir and *I*(*t*) is the input signal (Equation 1). *z_out_*(*t*) denotes the activity of the readout unit together with the output weights, *w_out_* (Equation 2). In our notation, *w_ij_* denotes the weight from neuron *j* to *i*, and so *w_out,i_* means the weight from *i^th^* unit in the reservoir to the readout unit. *ϕ* is the activation function given by:

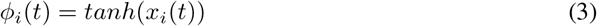

We use recursive least squares (RLS) to adjust the output weights, *w_out_* during training ([11]). The algorithm and the update rules are given by:

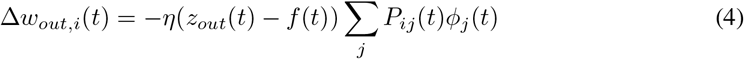

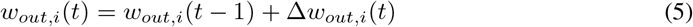

Here, *η* is the learning rate, *f*(*t*) is the target function, and the term ∑_*j*_ *P_ij_*(*t*)*ϕ_j_*(*t*) acts as a regularizer where *P* is the inverse cross-correlation matrix of the network firing rates. For details on setting hyperparameters, see Appendix A.

### 2.2 Adjusting reservoir dynamics

During training, the output weights, *w_out_*, are optimized using RLS based on the instantaneous difference between the output, *z_out_*(*t*), and the target function, *f*(*t*). Here, we use the reservoir to autoencode the input signal, thus *f*(*t*) = *I*(*t*). The instantaneous difference gives rise to an error term, *E*(*t*), calculated as:

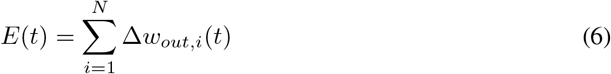

### 2.3 Obtaining the error signal

After training, the output weights, *w_out_*, are frozen. The test pattern is fed to the network via the input, *I*(*t*), and the network is iterated to obtain the error, *E*(*t*), over the duration of the test signal. The error, *E*(*t*), is computed as the difference between the test signal and the network output (Equation 6). The error may vary depending on the similarity of a given test signal to the learned time series. The error can be used directly to differentiate patterns, or as input to a classifier.

### 2.4 Classification of the error signal

The error, *E*(*t*), is used as input to a logistic regression classifier with L2 regularization. The classifier is trained using stratified 10-fold cross-validation. The same classifier and training procedure was used in comparing the different approaches. Naive Bayes and the multilayer perceptron (MLP) are directly used as classifiers, again with stratified 10-fold cross-validation. Accuracy and area under the curve (AUC) are computed as a measure of classification performance. See Appendix B for details on the specifics of the processor used for analysis and comparisons.

### 2.5 Neural recordings

#### 2.5.1 OFC data

Neural recordings were obtained from the macaque OFC using a custom designed 128-channel micro-ECOG array (NeuroNexus), with coverage including anterior and posterior subregions (areas 11/13). An experimental task was designed to understand how OFC helps in updating behavioral preferences in the face of unexpected outcomes. The details of the behavioral task design and data pre-processing can be seen in Appendix C.

#### 2.5.2 SCC data

8-channel neural recordings (local field potentials) were also obtained from the subcallosal cingulate (SCC) region of human patients who have treatment-resistant depression, undergoing deep brain stimulation in the OR (4 channels per hemisphere). The details of the recording protocol ((Figure 4A) and data pre-processing can be seen in Appendix D.

## 3 Results

### 3.1 Differentiating time-series patterns from different chaotic data recurrent neural networks (RNNs)

First, we trained TRAKR on multi-unit time series generated by chaotic data recurrent neural networks (RNNs) (Figure 2). Readout weights connected to the reservoir were fitted to the multi-unit signal using recursive least squares (RLS; see subsection 2.1). TRAKR is able to detect deviations from the trained pattern in real-time (Figure 2A). The network was trained on single-trial data from 30 units in a chaotic data RNN (Figure 2A, *Train* panel). With readout weights frozen, a *Test* signal was fed to the reservoir which contained a square-wave artifact (see section 2 for further details). The network *Output, z_out_*(*t*), and the *Error* signal, *E*(*t*), are depicted in the third and fourth panels from the top, respectively (Figure 2A). The network correctly detects the deviation of the *Test* pattern from the *Train* signal in real-time, evident as a sharp increase in the *Error* signal around the time point when the artifact is introduced Figure 2A, *Error* panel). The rest of the *Test* pattern corresponds to the *Train* pattern and, thus, correctly yields no further error spike.

**Figure 2:**
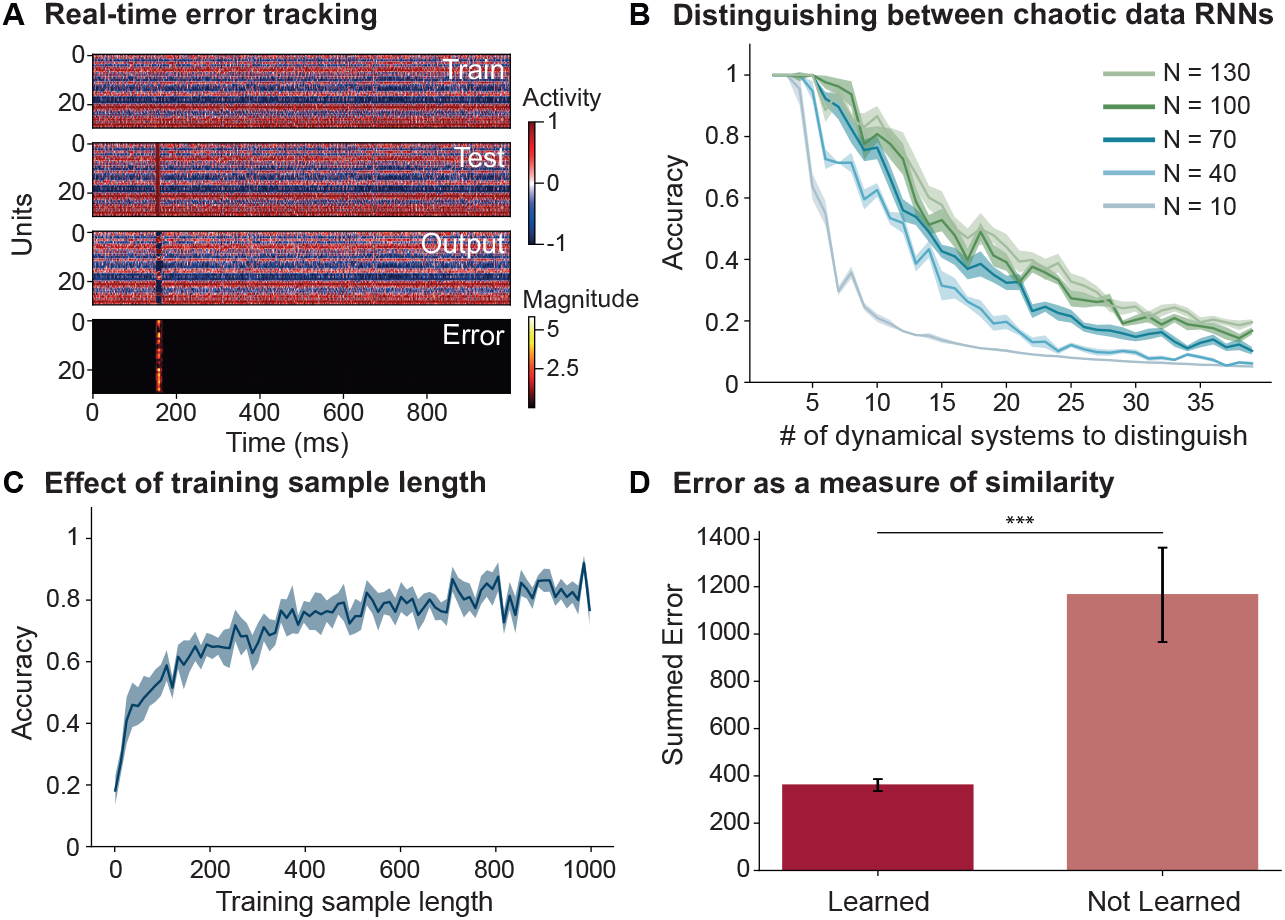
A) Error tracking of a square wave artifact in the test signal. Train signal (*w_out_* plastic), test signal (*w_out_* aplastic), TRAKR output and error signal shown from top to bottom. Error shows a spike at the time of the artifact in the test signal. B) TRAKR shows above-chance accuracy in classifying between upto 40 patterns from chaotic data RNNs. The five curves correspond to different number of units in the reservoir. C) TRAKR accuracy of classifying between 10 different patterns from chaotic data RNNs increases as a function of the length of the 10 patterns used for training (chance-level at 10%). D) The summed errors can differentiate between learned and new chaotic dynamical systems (∗ ∗ ∗ : *p* < 0.001).

We then trained TRAKR on data from different chaotic data recurrent neural networks (RNNs) to probe the capacity of the network with regards to differentiating between complex time-series patterns (Figure 2B). We trained TRAKR on single examples of data (1000 samples each) from chaotic RNNs. We successively trained TRAKR on examples from different chaotic data RNNs, considering the M-way classification problem with *M* ∈ [1, 40]. Considering the 10-way classification problem, for example, TRAKR was successively trained on single example traces from 10 different chaotic RNNs. Then we froze the readout weights, fed test examples from each of the 10 chaotic RNNs into the network (using randomly generated patterns from the chaotic RNNs that had not been used in training) and obtained an error signal for every test pattern. We then investigated whether it was possible to distinguish the 10 chaotic dynamical systems based on the error signals (see subsection 2.4 for further details). We found that it was possible to distinguish between the 10 chaotic dynamical systems based on the error generated by TRAKR with 86% accuracy (10-fold cross-validated, with 130 units in the network). As further patterns from different chaotic data RNNs are added, performance decreases gradually. TRAKR is still able to distinguish 20 chaotic dynamical systems with ≈ 45% accuracy (chance-level at 5%). Performance levels out at around 40 dynamical systems, but still remains above chance (20%, with chance-level at 2.5%). We also investigated the effect of different network sizes (Figure 2B), and found that performance improves with higher reservoir size, levelling out at 130 units.

We also investigated the effect of training sample length on TRAKR performance (Figure 2C). We focused on the 10-way classification case and measured accuracy when the training sample length for each pattern was varied from 1 to 1000 samples. We found that accuracy monotonically increases with training sample length, plateauing at ≈ 600 samples (10-fold cross-validated).

We also investigated if the summed error was enough to differentiate between learned and new patterns from chaotic dynamical systems not used in training (Figure 2D). We again took single example traces from 10 different chaotic data RNNs and successively trained TRAKR on them. We then obtained test traces (not used during training) from the 10 chaotic data RNNs which had been used to generate the training data, as well as from another set of 10 chaotic data RNNs which had previously not been used. We obtained error traces for all of the test patterns after passing through TRAKR, and summed the error. We found that the summed error was enough to differentiate between learned and new chaotic dynamical systems (*p* < 0.001, *std.dev*. across the 10 chaotic data RNNs in each category). Overall, this shows that TRAKR-derived error signal from single-trial examples with a sample length of ≈ 600 samples is sufficient to distinguish patterns from different chaotic dynamical systems with high accuracy.

### 3.2 Comparing TRAKR performance on a benchmark time-series dataset: permuted sequential MNIST

We applied TRAKR to the problem of classifying ten digits from permuted sequential MNIST, a benchmark time-series dataset ([19, 18]), and compared its performance to other commonly used approaches.

We used a dataset of 1000 permuted sequential MNIST digits (including 100 samples for each digit (0-9)) to compare TRAKR with other approaches commonly employed in time-series classification (Figure 3). Every image of a digit (28 x 28 pixels) was flattened into a vector, and the order of the pixels was randomly permuted. For training, we randomly drew a sample from one of the digits and trained TRAKR on that single example of a particular digit (see subsection 2.1 for training details). After that, we froze the readout weights and obtained error traces, *E*(*t*), for all the test samples from each of the 10 digits. The error traces were used to train a classifier (see subsection 2.4 for more details). We repeated this procedure for all digits in the dataset to arrive at the final averaged TRAKR classification performance (Figure 3A).

**Figure 3:**
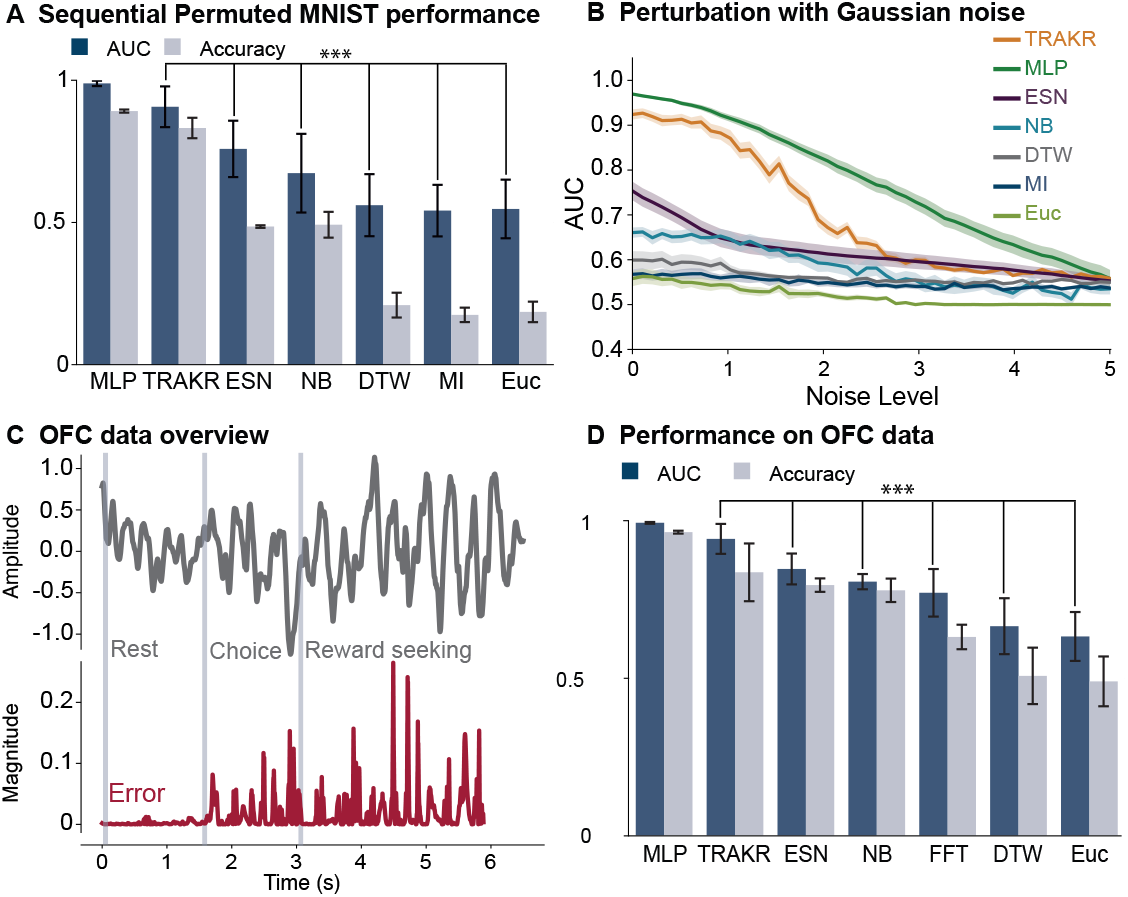
Classification performance on the permuted sequential MNIST and macaque OFC datasets. A) TRAKR performs on par with MLP, while outperforming all other methods in classifying the permuted sequential MNIST digits (92%AUC; ∗ ∗ ∗ : *p* < 0.001, Bonferroni-corrected; chance-level at 10%). MLP: Multilayer perceptron (as in [10]); ESN: echo-state network (twiESN, as in [10]); NB: Naive Bayes; DTW: Dynamic Time Warping (as in [27]); MI: Mutual Information; Euc: Euclidean distance metric. B) Classification performance with increasing amounts of noise added to the digits. TRAKR performance declines smoothly with noise level, while still outperforming most other approaches in classifying noisy digits. Chance level is at 50%. C) Grey - example neural time series from a single trial, with three behaviorally relevant epochs (rest, choice and reward seeking periods). Red - associated error trace when trained on the rest period of a different reference trial and tested on this entire trial. D) TRAKR performs on par with MLP, while outperforming all other methods in classifying the different neural epochs (∗ ∗ ∗ : *p* < 0.001, Bonferroni-corrected; chance-level at 33%). FFT: Fast Fourier Transform.

We found that TRAKR achieves a performance of *AUC* = 92% on the permuted sequential MNIST dataset (Figure 3A). We compared our approach with other commonly used methods for the classification of time series, such as deep supervised neural networks (or multilayer perceptrons, MLP; as implemented in [10]), another recent echo-state (or reservoir) network-based approach (twiESN; as implemented in [10]), Dynamic Time Warping (DTW, as implemented in [27]), Euclidean distance (Euc), Mutual information (MI), and a naive Bayes classifier (NB). We used the same classifier to compare performance to other distance metrics as well as DTW (see subsection 2.1 for details). TRAKR performed significantly better than ESN, NB, DTW, MI and Euc (*p* < 0.001, 10-fold crossvalidated), and on par with MLP (which employed multiple batches of training data and several successive rounds of optimization).

We also compared performance when the training data was perturbed with Gaussian noise (Figure 3B, see also Appendix E). For this purpose, we added Gaussian noise of varying levels. We found that TRAKR again performed better than other approaches such as ESN, NB, DTW, MI and Euc, and comparably to MLP, though MLP performed best. TRAKR performance decayed gradually until reaching chance-level at a noise of *σ* ≈ 3.

### 3.3 Comparing performance on biological data from the macaque orbitofrontal cortex (OFC)

The OFC is a higher-order brain region involved in encoding affective expectations ([31, 32]). We used neural data from the OFC recorded during a behavioral task designed to reveal how such expectations were encoded in the OFC of primates (see subsubsection 2.5.1 and Appendix C for more details). The goal here was to determine whether TRAKR could be used to reliably distinguish complex neural time-series patterns based on single-trial data, and how it compared to other approaches.

A recording from a single electrode in a single trial is shown spanning three different behaviorally relevant epochs (*rest, choice* and *reward seeking* periods) (Figure 3C, grey trace). The goal was to distinguish the neural patterns in these three epochs. Importantly, it was not known a priori whether there was enough information contained in the OFC recordings to allow for differentiating the three epochs; a high-performing classifier can reveal what kind of information is encoded in an area such as OFC, in the brain.

We again compared TRAKR with all other approaches on the problem of distinguishing the neural signals from these three epochs (Figure 3D). We trained TRAKR on the neural time series corresponding to rest period from a particular trial, and used the other complete trials as test signals to obtain errors, as before. The TRAKR-derived error-trace is depicted for the above example (Figure 3C, red trace). The magnitude of the error during the initial rest period, which TRAKR was trained on, is relatively lower, compared to the error for the other two periods. A classifier was trained on the error traces, as before. In addition to all the methods introduced before, we also calculated the Fast Fourier transform (FFT) of the signals and obtained the magnitude (power) in the *α* (0 – 12*Hz*), *β* (13 – 35*Hz*), and *γ* (36 – 80*Hz*) bands within the 3 epochs and included this in the comparison.

We found that TRAKR outperformed all the other approaches with the exception of MLP (*AUC* = 94%; *p* < 0.001; Figure 3D). TRAKR was able to distinguish the three epochs with high accuracy based on single-trial data, showing that there is enough information in the OFC to differentiate these patterns. Notably, based on the performance of lower performing approaches, such as DTW or Euc, one might have (wrongly) concluded that the OFC did not encode information about other task variables beyond the rest period.

Overall, this shows that TRAKR can be used as a reliable tool to differentiate time-series patterns in biological data and probe what kind of information is encoded in different brain regions.

### 3.4 Distinguishing neural time-series recorded from the human subcallosal cingulate cortex (SCC)

Lastly, we applied TRAKR to an 8-channel neural time-series dataset recorded from the SCC of human patients (Figure 4). The overall goal of the recording protocol in the OR was to find neural signatures (or biomarkers) of behavioral effects observed in response to intraoperative brain stimulation ([34]; see subsubsection 2.5.2 for more details). For a particular individual, 8-channel recordings were obtained after stimulation on each contact (Figure 4A). We trained TRAKR on 8-channel recordings after stimulation at one contact (*E*1, in particular, as depicted in Figure 4A), and used 8-channel recordings after stimulation at all other contacts as test samples, thus using TRAKR to examine differences in the post-stimulation effects across all electrodes. We obtained activations for all the units in the reservoir for the test samples, and projected them into a common space spanned by the first 3 principal components (PCs) of the activations. We plotted the mean activity of TRAKR’s reservoir in this space (Figure 4C), and compared it to the 3-D representation of the 8-channel recordings (after stimulation at different contacts) by fitting the PCA model directly to the neural data (Figure 4B).

**Figure 4:**
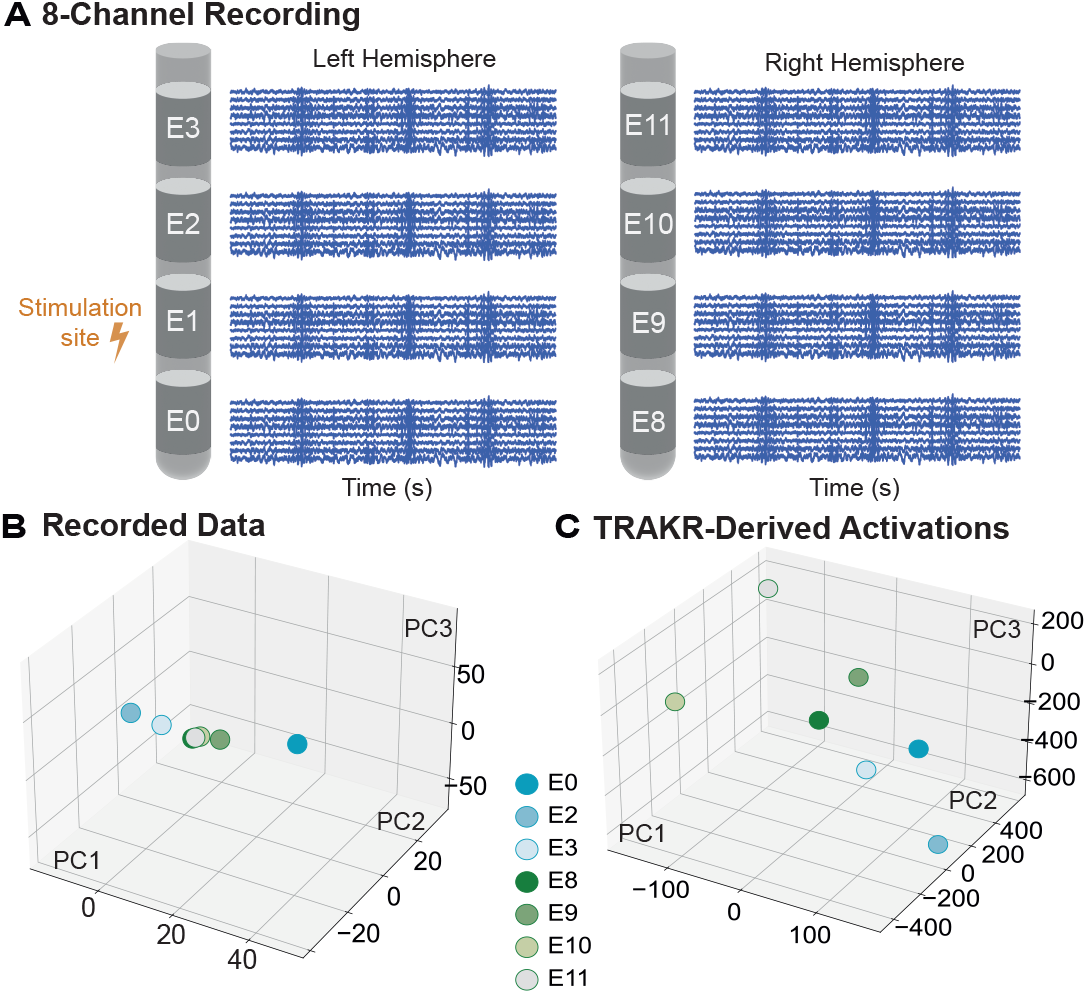
A) 8-channel recording protocol (see subsubsection 2.5.2 and Appendix D for detailed description). Contacts in the left hemisphere: *E*0 to *E*3; in the right hemisphere: *E*8 *to E*11. B) 3-D representations of the 8-channel recordings after stimulation on different contacts. Effect of stimulation not noticeable. C) 8-channel recordings after stimulation on different contacts projected into the space spanned by the first three principal components of TRAKR activations. Recordings after stimulation on left contacts are separable from those after stimulation on the right. Legend shows the stimulation contact after which 8-channel recording was obtained.

We found that TRAKR-derived activations allow for better detection of the effect of stimulation: 8-channel recordings after stimulation on left hemisphere contacts (*E*0 to *E*3) are more easily separable from those after stimulation on the right (E8 to E11) with TRAKR derived representations (Figure 4C). Comparatively, the effect is not easily noticeable based on the representations obtained from performing dimensionality reduction on the 8-channel recordings directly (Figure 4B).

## 4 Discussion

We have shown that TRAKR can be used for accurate time-series classification based on singletrial data. It can reliably distinguish a high number of patterns from chaotic data RNNs with high accuracy. TRAKR outperforms other commonly used approaches (such as Euclidean distance, Mutual information, or DTW) on a benchmark dataset, permuted sequential MNIST, and on biological data from the neuroscience domain. It performs on par with supervised approaches based on deep neural networks (MLPs) trained with gradient descent on multiple batches of training data.

TRAKR also performs better than another recent reservoir-based approach (twiESN, as in [10]) on both permuted sequential MNIST and on biological data. This shows that our approach based on the error trace is key to the performance we observe here. We also show how imaging the activations inside of TRAKR’s reservoir can be used to visually distinguish neural time-series patterns in a lowdimensional space. The separation achieved with TRAKR is better than that achieved by using PCA directly on the neural recordings. This suggests TRAKR can be used in conjunction with PCA to improve clustering results in unsupervised approaches, for example.

One current limitation is that we employ a classifier on top of the error traces to perform the classification here. Nonetheless, we show that the summed error is enough to differentiate between learned and new chaotic dynamical systems (Figure 2D). Thus, in future work, we aim to add an additional layer with sigmoid readouts so as to allow for the classification result to be directly read out from the TRAKR reservoir.

This work contributes a new approach to the classification of time-series data that offers high accuracy results on single examples of data from the neuroscience domain. As such, we do not anticipate any direct ethical or societal impact. Over the longer term, we believe our tool can have impact on related research communities such as neuroscience and clinical research, with societal impact depending on the development of these fields.

## 5 Conclusion

There is a need for and strong interest in tools for the analysis of time-series data ([5]). We show that TRAKR is a fast, accurate and robust tool for the classification of time-series patterns. Through its ease of use and low computational overhead, it is particularly suited for real-time applications where accurate decisions need to be made quickly and signal degradation or other artifacts necessitate frequent re-calibration. TRAKR can also be used to distinguish neural time-series patterns in the brain based on single-trial data, generating new hypotheses about information processing in neural circuits.

## Acknowledgments and Disclosure of Funding

This work was funded by NIH 1*R*01*EB*028166 – 01 (Dr. Rajan), NSF FOUNDATIONS Grant 1926800 (Dr. Rajan), NIH *UH*3*NS*103550 (Dr. Mayberg), Pew Biomedical Scholars Program supported by the Pew Charitable Trusts (Dr. Rich) and NARSAD Young Investigator Grant from the Brain & Behavior Research Foundation (Dr. Rich). We also thank Aster Perkins for neural data collection.

## A TRAKR Hyperparameters

The recurrent weights *J_ij_* are weights from unit *j* to *i*. The recurrent weights are initially chosen independently and randomly from a Gaussian distribution with mean of 0 and variance given by *g*^2^ /*N*. The input weights *w_in_* are also chosen independently and randomly from the standard normal distribution.

An integration time constant *τ* = 1*ms* is used. We use gain *g* = 1.4 for all the networks.

The matrix *P* is not explicitly calculated but updated as follows:

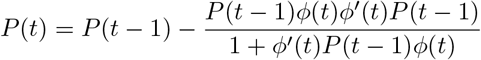

The learning rate *η*, is given by 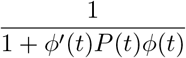.

The number of units used in the reservoir is generally *N* = 100.

## B Processor Specifics

MacBook Pro CPU was used for all the comparisons. The specifications include a 2.3 GHz 8-core Intel Core i9 processor with 32 GB RAM.

## C Behavioral task design and data pre-processing - OFC data

The OFC data was obtained for 2 different macaques, and data from one is analyzed and shown in the submission. The data was collected at a medical institution.

During preliminary training, the monkey learned to associate unique stimuli (natural images) with rewards of different values. Rewards were small volumes of sucrose or quinine solutions, and values were manipulated by varying their respective concentrations.

During the task, the monkey initiated a trial by contacting a touch-sensitive bar and holding gaze on a central location. On each trial, either one or two images were presented, and the monkey selected one by shifting gaze to it and releasing the bar. At this point, a small amount of fluid was delivered, and then a neutral cue appeared (identical across all trials) indicating the start of a 5s response period where the macaque could touch the bar to briefly activate the fluid pump. By generating repeated responses, it could collect as much of the available reward as desired. There were two types of these trials. Match (mismatch) trials were those where the initial image accurately (did not accurately) signal the type of reward delivered on that trial. Behavioral performance and neural time series were recorded in 11 task sessions across 35 days. Each trial was approximately 6.5s long, including different behaviorally relevant epochs and cues. The macaque performed approximately 550 trials within each task session (*mean* ± *sd* : 562 ± 72). Of note, 80% of the trials were match trials within each task session.

ECoG data were acquired by a neural processing system (Ripple) at 30*kHz* and then resampled at 1*kHz*. The 128-channel data were first z-score normalized. Second-order butterworth bandstop IIR filters were used to remove 60*Hz* line noise and harmonics from the signal. We also used second-order Savitzky-Golay filters of window length 99 to smooth the data and remove very high frequency juice pump artifacts (> 150*Hz*). For most of the analysis here, we used the average of the 128-channel time series as an input to TRAKR.

## D Recording protocol and data pre-processing - SCC data

The data was collected at a medical institution, where eight patients with treatment-resistant depression underwent deep brain stimulation surgery in the operating room. IRB approval and consent was obtained from all the patients before the surgical procedure. The exploratory analysis shown in this paper is from one of these patients. The data was anonymized and did not contain personally identifiable information.

These neural recordings were obtained using the Alpha Omega system in the OR. The recordings were performed under a stimulation protocol such that each contact was electrically stimulated for 3 minutes followed by 1 minute of recording without stimulation. For all such 1 minute recordings, 8-channel neural time-series data (local field potentials) were obtained from the SCC region. That gives 8 sets of 8-channel recordings (after stimulation on 8 contacts).

Data were originally acquired at 22 *kHz* and then resampled at 1 *kHz*. Data were notch-filtered around line noise, its harmonics and stimulation frequencies to remove the artifacts from the physiological signal. Then, the data were band-pass filtered between 0.5 and 80*Hz*. Data were standardized and common average referenced.

## E Noise addition to permuted sequential MNIST digits

For measuring noise robustness, we added random independent Gaussian noise to the training digits (*μ* = 0 and varying standard deviation (*σ*)).

